# Synthetic Turing patterns in engineered microbial consortia

**DOI:** 10.1101/2020.06.15.153205

**Authors:** Salva Duran-Nebreda, Jordi Pla, Blai Vidiella, Jordi Piñero, Nuria Conde, Ricard Solé

## Abstract

Multicellular entities are characterized by exquisite spatial patterns, intimately related to the functions they perform. Oftentimes these patterns emerge as periodic structures with a well-defined characteristic scale. A candidate mechanism to explain their origins was early introduced by Alan Turing through the interaction and diffusion of two so called *morphogens*. Unfortunately, most available evidence for Turing patterns in biology is usually obscured by the tangled nature of regulatory phenomena, making difficult to validate Turing’s proposal in developmental processes. Here we follow a different approach, by designing synthetic genetic circuits in engineered *E. coli* strains that implement the essential activator-inhibitor motif (AIM) using a two-cell consortium. The two diffusible compartments are one cell type (activator, small-diffusion component) and a small signal molecule (a homoserine lactone, acting as the fast-diffusing inhibitor). Using both experimental results, we show that the AIM is capable of generating diffusion-induced instabilities leading to regular spatial patterns. The artificial construction taken here can help validate developmental theories and identify universal properties underpinning biological pattern formation. The implications of the work for the area of synthetic developmental biology are outlined.

## I. INTRODUCTION

The rise of multicellular life forms defines one of the major transitions in evolution, and required novel ways of organisation grounded in the cooperative interactions among single cells within supracellular assemblies. The emergence of developmental processes and gene regulatory networks provided a flexible and adaptive source of morphological diversity, facilitated by a number of physico-chemical generative mechanisms [1–3]. These mechanisms allow the emergence of long-range order out of locally interacting cells, and they often rely on signalling molecules diffusing in space [2, 4], particularly at early stages of development [1, 5–8].

An elegant mechanism for the emergence of long-range order out of short-range interactions was formulated by Alan Turing [9, 10]. In his seminal paper *On the chemical basis of morphogenesis*, Turing suggested that a system composed of two diffusing and interacting molecules (an activator and an inhibitor) could explain how an initially homogeneous state could lead to regular macroscopic structures by means of amplification phenomenon. Specifically, Turing showed that the interaction between a pair of molecular species displaying diffusion can, under some mathematical conditions, trigger *diffusion-driven instabilities* [9, 11–14]. The basic motif (the activator-inhibitor motif, AIM) linking the two morphogens is depicted in (see figure 1a). Along with the feedbacks connecting activator (*A*) and inhibitor (*I*) molecules, a marked asymmetry exists between their diffusion rates *D*_1_ and *D*_*i*_, respectively: the activator has a much shorter diffusion range than the inhibitor molecule (*D*_*i*_ ≫*D*_*a*_). Under these conditions, it was predicted that ordered spatial patterns would emerge with a characteristic spatial scale much larger than the cellular one.

**FIG. 1:**
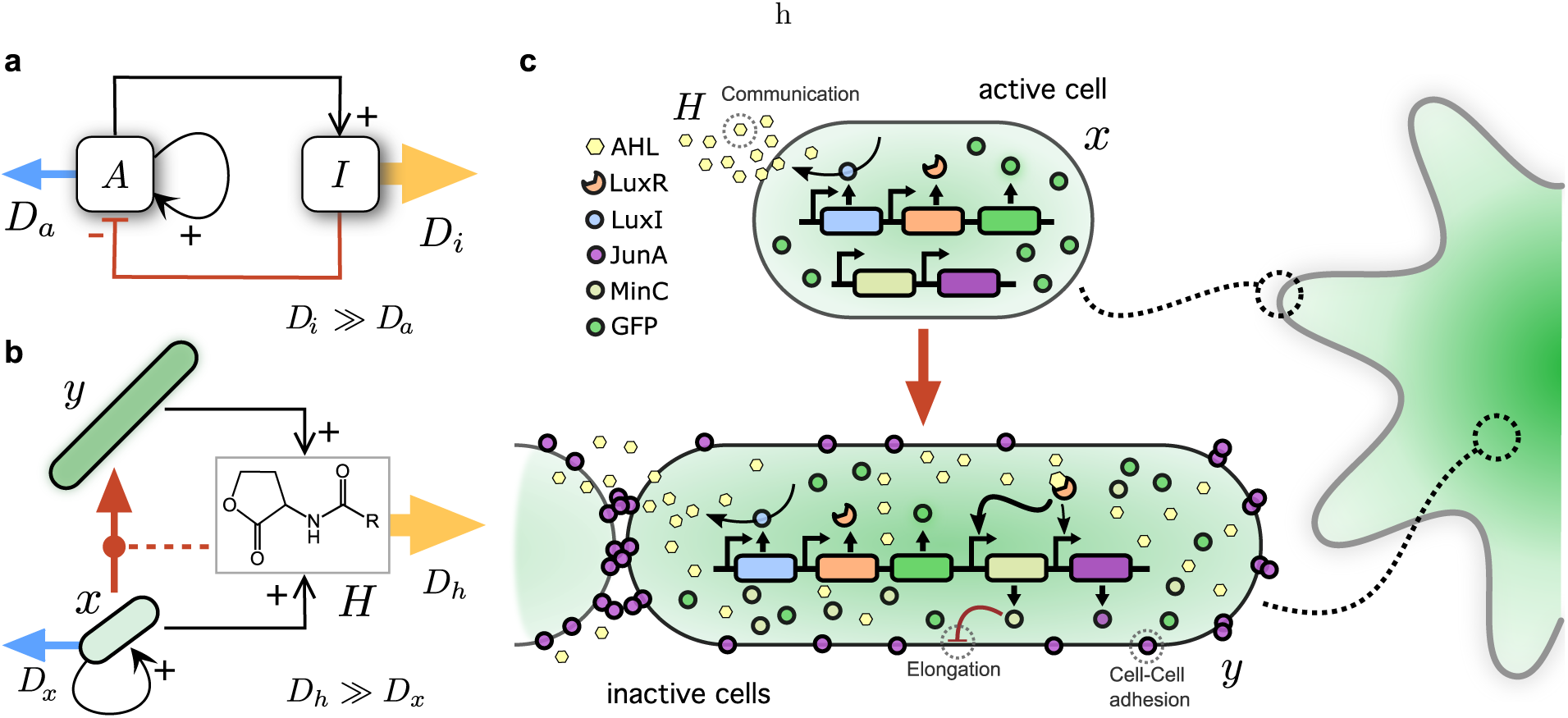
Engineering Turing patterns. In (**a**) the standard mechanism leading to diffusion-driven instabilities is shown. Here *A* and *I* indicate the two minimal morphogens i. e. the activator and inhibitor, respectively. The arrow thickness indicates different diffusion rates associated with each morphogen. The standard motif summarized here (**b**) the corresponding diagram for our engineered strains is shown. Here the initial cell population (*x*) can differentiate, under the presence of *H*, in an elongated phenotype that does not replicate (but also produces *H*) nor diffuse. The *H* molecule is an inhibitor anting both directly on *x* and indirectly through its effects on differentiation. A more microscopic picture of this network motif is shown in (c) where the genes involved in the full engineered system.

Turing-like patterns have come to be regarded as a paradigmatic example of simple systems yielding complex features and have been successfully found in both physical and chemical systems [15–18], and strong evidence has been found in biology. This includes skin pigmentation in animals [19–26], primordia of skeletal elements [27–30], palatal ridges [31], teeth formation [32], establishment of hair follicles [33] and ecological systems [34–39].

Nonetheless, experimental evidence for Turing instabilities in developmental biology has been debated, with most advances based on theoretical models and mathematical extensions of the theory [40–42]. Only recently, some experiments have begun to unravel the picture of Turing-type mechanisms in the formation of skeletal limb primordia [29, 30], suggesting that more morphogens and more complicated interaction maps might be involved than originally thought. However, an inevitable result of metazoan regulatory networks is that the variables involved are many more than those associated to the simple Turing mechanism, thus obscuring the logic of the underlying process.

Another possibility for inquiry into pattern formation is given by synthetic biology [43, 44] which tries to artificially construct -or reconstruct-complex functions and features by splicing together coding and regulatory sequences that do not naturally coexist [45–47]. Such approach has enabled researchers to mimic gradient-based positional information developmental systems [48–50], however self-organized systems -characterized by simple interacting units and horizontal exchanges of information like in Turing’s mechanism-have proven to be more elusive to engineer [51–53].

Successful experimental implementations of Turing-like instabilities include a recent engineering of spatial patterns inspired in the short-range inhibitor, long-range activator scheme (figure 1a) engineered on a synthetic mammalian system. Using the Nodal-Lefty signaling system, it was actually possible to illuminate the nature of interactions between this pair of signals [54]. One particularly nice outcome of the model was that, by manipulating diffusion rates and making they equal, no pattern emerged. This is consistent with the conditions required for Turing patterns to emerge. Some strategies use stochastic Turing theory [55]. In that case, demographic noise can induce persistent spatial pattern formation that is reflected by the presence of (statistical) spatial correlations.

Here we report a novel way of designing a synthetic pattern-forming system using bacteria that involves communication, filamentous growth, adhesion and growth inhibition. As shown below, our engineered cells are able to create spatially regular structures consistent with those created by Turing-type mechanisms. This research provides an important milestone in the establishment of mutual feedbacks between experimental embryology, modeling of pattern formation and synthetic strategies to reconstruct putative mechanisms and interactions. By using bacterial growth on Petri dishes, we take advantage of the pattern-forming capabilities of microorganisms [56–59].

However, our model differs from other models of bacterial colony growth displaying spatial patterning in one fundamental point. Instead to relying in a limited available resource that is depleted along the growth of the microbial population, resources are in excess to prevent that mechanism from influencing our systems. Under such conditions, the reaction-diffusion motif defined below is the only one responsible for the emergence of regular patterns. Additionally, the role of “activator” is played by cells (thus diffusing at low rates) whereas inhibition is engineered using a small (homoserine lactone, AHL) diffusible molecule. This approach was shown to properly define a pattern-forming instability in a computational model of engineered populations involving chemotaxis [60]. The interaction between both is mediated by a third partner that allows completing the basic logic required for Turing instabilities to occur. The differentiated compartment *y* exhibits elongated cells that attach to each other by adhesion molecules. As a result they will tend to accumulate in domains where *H* concentrations are higher and cell replication rates small (as expected in the activator-inhibitor picture). On the other hand, as growth centers emerge on the periphery of the population boundaries, they will keep growing but experience inhibition of nearest centers due to the effects of *H*. If such spatial inhibition operates under a Turing-like process, we should expect growing centres to inhibit each other and remain separated by a characteristic length scale.

## II. RESULTS

### 1. Synthetic pattern-generator design

The logic of our designed circuit is summarized in Figure 1b-c. In figure 1a, we display the standard Turing motif where two diffusible molecular populations are indicated by *A* and *B* and interact as depicted: the activator grows by means to an autocatalytic process, while positively helps increasing the inhibitor populations. The later instead acts inhibiting the production of the former. Diffusion of *A* and *I* are short- and Long-ranged, respectively. This basic topological structure of the network motif is preserved in our construct (figure 1b) but is mediated by a microbial consortium where two different cell types are present, as a result of a differentiation process *x → y*. Their populations are indicated by *x* and *y*. This consortium involves a growing population (*x*) that expands while it produces the inhibitor molecule *H* and is inhibited back by *H*. Additionally, *x* differentiates into *y* (a non-growing population of elongated cells) under the presence of *H*.

The overall balance of this design is that cells cause increase the concentration of the inhibitor molecule, which diffuses much faster than the dispersal of cells from the *x* compartment. In figure 1c we also summarize the microscopic interactions happening in our experimental implementation, indicating the basic genetic parts used in the design (to be detailed below).

### 2. Non-homogeneous spatial distributions of cells arise at the intersection between adhesion, cell signaling and filamentous growth

In this study we have assessed the impact of four genes (Fig. 2a) in the colony growth of *E. coli* UT5600 strain, which typically develops into uniform circular colonies as time progresses. The genes synthetically introduced in the model organism comprise different effects in cell morphology, growth rates, cell-cell adhesion and cell-cell signaling through quorum sensing.

**FIG. 2:**
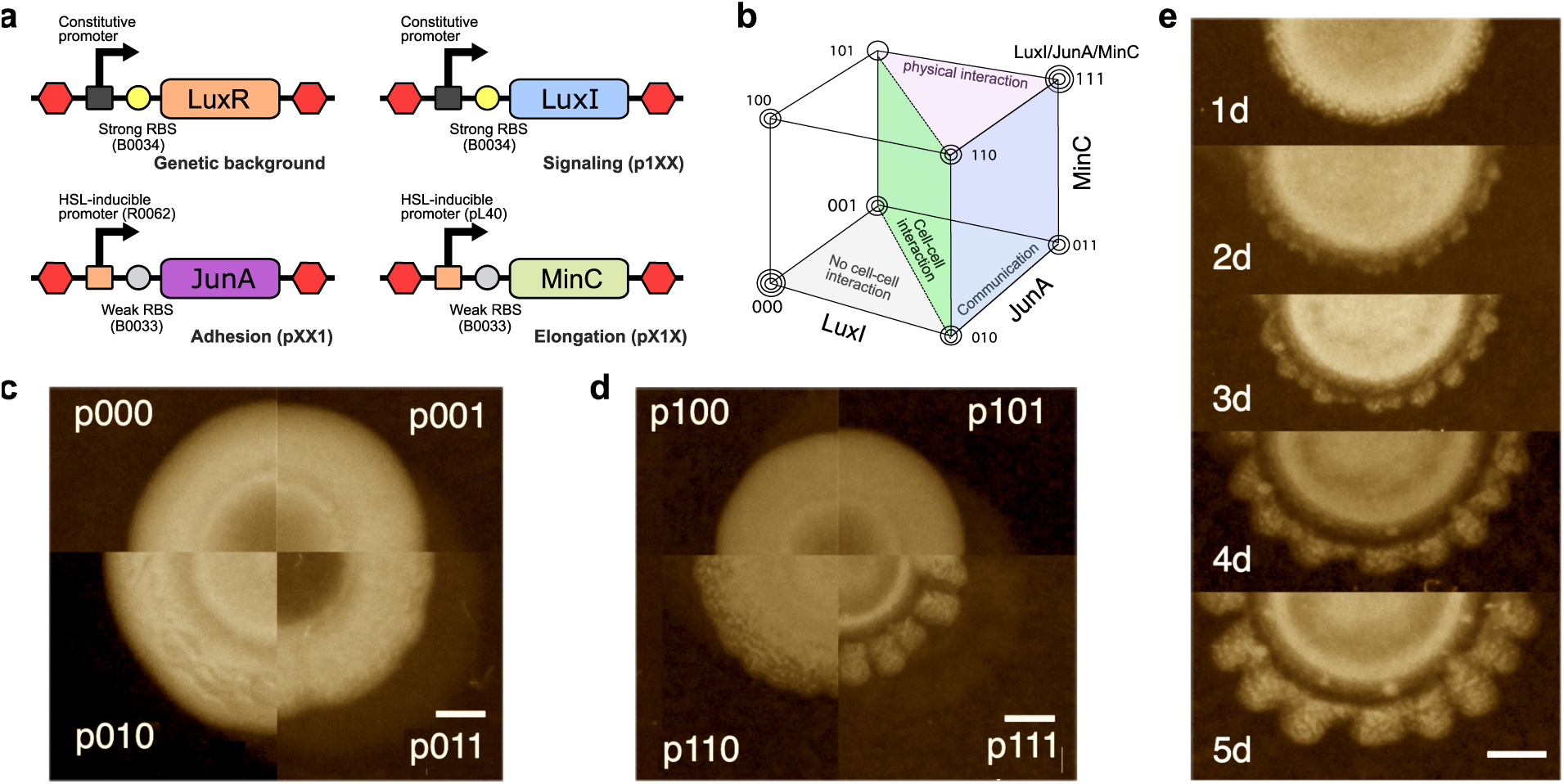
Regular patterns are only achieved at the intersection between communication, adhesion and elongation. (**a**) Basic scheme of the constructs used in this work. Receiver gene (*LuxR*) was present in all tested strains while the other three were introduced in all possible combinations. Notation goes as follows, each bit indicates the presence or absence of a particular feature, from left to right: signaling, elongation and adhesion. In the morphospace resulting from the combination of the three possible elements (**b**) it can be seen different regions: the separation between individual behaviour to cell-cell interactions (green surface), physical interaction (purple), and community communication/synchronization (blue). Patterns are only obtained when the three elements are present. (**c-d**) The eight strains evaluated in this work after 5 days of culture. The group on the right contains all combinations capable of cell-cell communication, while those on the left were exogenously supplemented with the signaling molecule. (**e**) Time series of p111 colony growth, daily progression. Scale bar in all pictures is 2 mm.

The first two items are given by *MinC*, a protein involved in *E. coli* segmentation [61, 62]. In particular, exogenous expression of this protein has been linked to the elongation of cells with the formation of filaments that can attain two orders of magnitude the typical cell length and entails a diminished biomass growth [63]. Cell-cell adhesion is introduced by a chimeric protein composed of the animal *JunA* coupled with an autotranslocator domain [64]. This chimeric sequence is able to target the membrane and homodimerize with equal proteins expressed by other cells, increasing the sedimentation rates in liquid cultures of bacteria. Finally, communication was incorporated by the expression of two components from quorum sensing system of *V. fisherii*, widely used by the synthetic biology community [48, 49, 51, 65, 66]. This is typically composed of a receptor protein (*LuxR*), able to enhance expression in specific promoter sequences in the presence of the ligand homoserine lactone (hereafter HSL), and the *LuxI* gene, able to synthesize the cognate molecule from preexisting substrates. This family of ligand molecules can passively diffuse across the cellular membranes, reaching high concentrations naturally when cellular density surpasses a threshold.

In order to explore the landscape of pattern formation capabilities of these genes implicated in cell morphology, adhesion and communication, all possible combinations of them were constructed. The space of possible combinations has been sketched in figure 2b, using the binary coordinates of the space (LuxI,JunA,MinC), where 1 indicates presence of the construct and 0 its absence, respectively. The eight possible combinations were tested and their impact in colony growth analyses. The spatial patterning of each colony after 5 days of growth are summarized in figure 2c-d. Those conditions lacking the ability to synthesize the signaling molecule were externally supplemented with HSL in the petri dish to ensure similar levels of *MinC* and *JunA* expression, but otherwise lack the spatial information given by the quorum sensing mechanism. Only when all three capabilities were included in the synthetic cells (i. e. at the (1, 1, 1) vertex of our binary space) spatial non-homogenous distributions of cell densities were created, in stark contrast to the other conditions were circular growth took place. With slight variations, each quadrant in the p111 pattern include five branches each.

### 3. Non-homogenous patterns are characterized by a dominant wavelength

As it can be seen in figure 2d, when all four genes are present the symmetry of the colony is broken in a regular fashion. The characteristic length scale is preserved along the colony growth process, as displayed in figure 2e, where different snapshots are shown. It is worth noticing that the regular branch pattern that stabilizes over time is preceded by a transient where some extra roughness is present. As expected from a Turing-like instability, there short-frequency irregularities are erased as the characteristic length is selected.

In order to characterize the higher order properties displayed by the growing structure, we analyzed the distribution of bacterial concentrations in a circumference centered at the (estimated) center of the colony. The goal is to quantitatively detect the presence of some characteristic length scale pointing to a regular, self-organizing pattern.

Since we lack a one-dimensional growing system (as it occurs with many models), the data set required first of all a transformation of bacterial density from Cartesian to polar coordinates (Fig. 3a-b). A Fast Fourier Transform (FFT) algorithm was used to calculate the power spectrum *P* (*f*) of the spatial data (Fig. 3b) scaling with population density. As expected, the one-dimensional mapping is affected by local fluctuations and asymmetries associated to the non-perfect circular shape, which tends to broaden *P* (*f*). Nevertheless, a well-defined peak is observed was found to display a characteristic wavelength correlation at approximately *λ ≈* 0.2 cm (Fig. 3c) consistent with the observed length scape of the pattern displayed by the colony (inset figure 3). For comparison, the Power spectrum for the p011 which does not reveal any periodic signal.

**FIG. 3:**
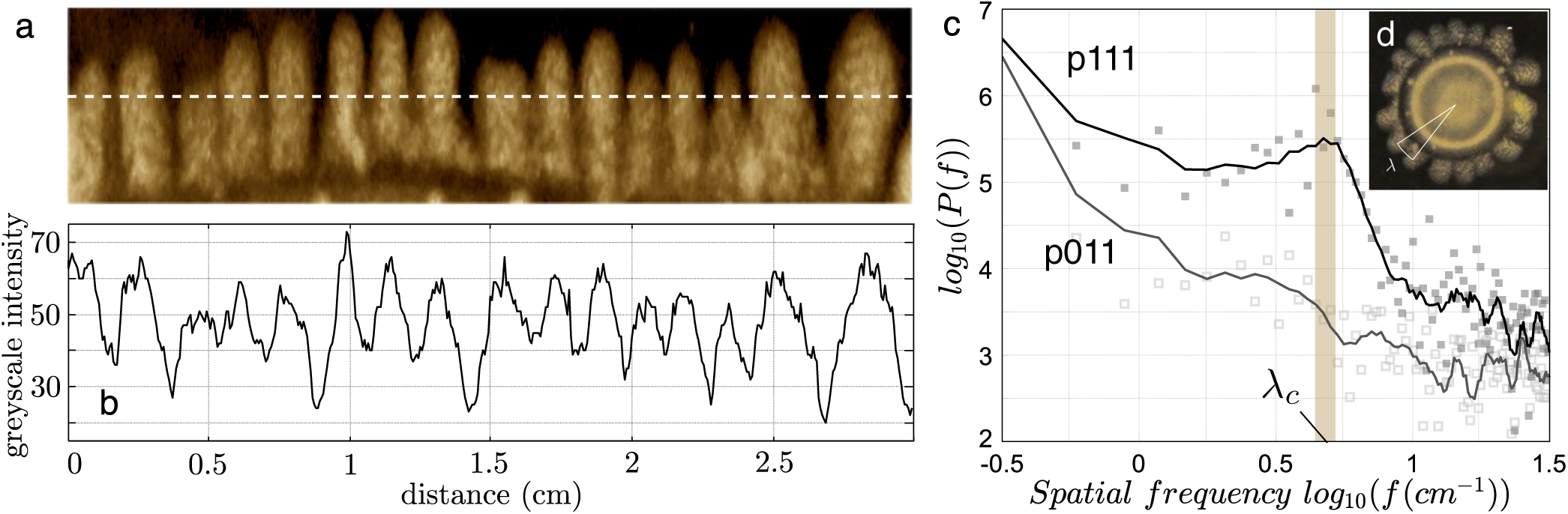
Characterization of the regular structures created by bacterial density. (**a**) Five-day colony profile after being transformed into polar coordinates. The intensity (a.u.) from the previous data is used as a surrogate of bacterial density close to the edge of the colony. A one-dimensional spatial set (**b**) is used to calculate the resulting power spectrum (**c**) showing the existence of a peak at a frequency value leading to a characteristic wavelength *λ ∼* 0.2 cm (brown shading) and consistent with the scale observed in mini petri-dishes (inset). Single colony assessment, for comparison we show the same analysis for a unstructured p011 colony (which lacks any characteristic scale).

### 4. Local and long-term patterns of branch primordia

In order to assess the micro-scale properties of the pattern formation process we captured the first stages of colony growth with bright field and fluorescence microscopy (Fig. 4a). In particular, we observed that the starting symmetry imposed by the circular droplet of cells is broken as soon as 24 hours when the whole set of genes is present. However, these primordia of the branching pattern do not have the regularity displayed by the colony at later stages, and are characterized by the formation of cohesive bundles of cells with the same orientation. Given that *E. coli* segmentation occurs perpendicular to the longest axis of the cell, the establishment of a collective orientation forcefully imposes growth in a preferred direction, which cells maintain in the following days. The synthetic cell adhesion features provided by Min along with the local directionality imposed by the E. Coli cell shapes can explain these features and might play some relevant role in the amplification mechanisms triggered by our synthetic Turing motif. In this context, the potential impact of cell phenotypes needs to be taken into account in the long-term shape changes.

**FIG. 4:**
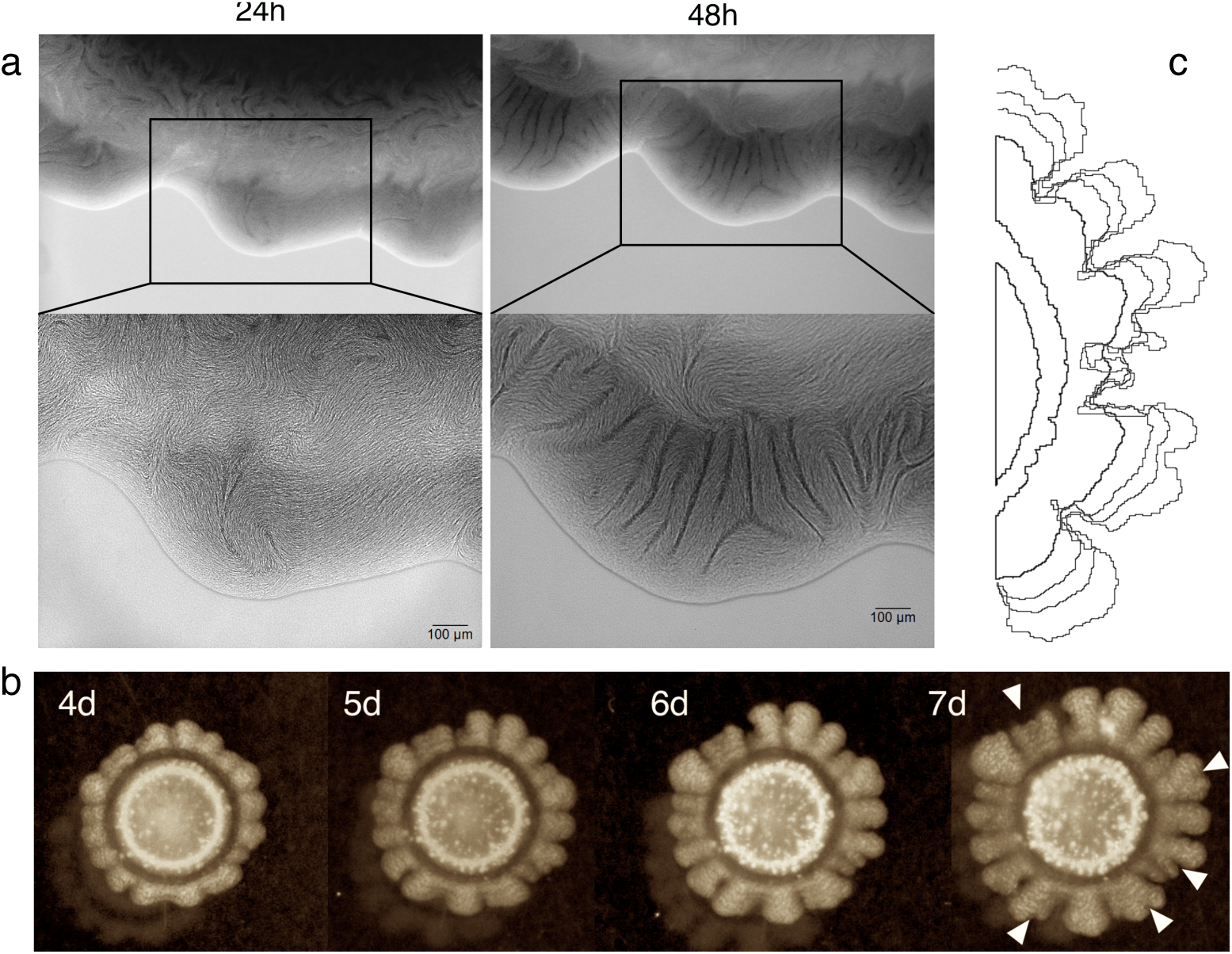
Small-scale and long-term pattern formation. (**a**) displays the same region of a p111 strain colony after 24 and 48 hours of growth in bright field microscopy. On the top, a wider region containing several branches, below the detail of cell bundles forming at the edge of the growing colony. (**b**) Time series of branch formation in older colonies. From the 4th day to the 7th we show the same colony and how once the zones of low density have been created they are not homogenized by the effects of diffusion. Marked with arrows are the new branching points: when domain width has surpassed a critical size, new troughs are created that branch the density. Scale bar represents 2 mm. (**c**) Overlaped edge colony detection with standard ImageJ libraries.

The long-term branching beyond the 5-day window used to determine the pattern-forming process reveals other qualitative properties that are consistent with the Turing instability process. In figure 4b-c a series of snapshots are used to see how the regular branching is maintained, but also how in the long run sprouts with a wide spacing experience new branching bifurcations (figure 4b, 7d snapshot). At this point, new instabilities might be taking place as a consequence of the Turing mechanism: since a characteristic *λ*_*c*_ is the only stable solution, structures with a larger wavelength will tend to split. This seems indeed to be the case. Some additional insight can be obtained looking at the changes in colony edges associated to this sequence (figure 4c) where we can appreciate how slight changes in edge curvature precede the emergence of branches.

### 5. *Lactone-induced expression of* MinC *and* JunA *inhibits growth of synthetic cells*

Following the previous examination we characterized the impact of the secreted quorum sensing molecule in the growth rate of cells. This was tested by monitoring bacterial density for different constructs at Abs_660_ in a liquid culture, with increasing amounts of externally introduced lactone. Figure 5 shows how the p011 strain (able to express *MinC* and *JunA* in the presence of inducer but unable to constitutively synthesize the signaling molecule) displays varying rates of growth and carrying capacities dependent on the concentration of the signal.

**FIG. 5:**
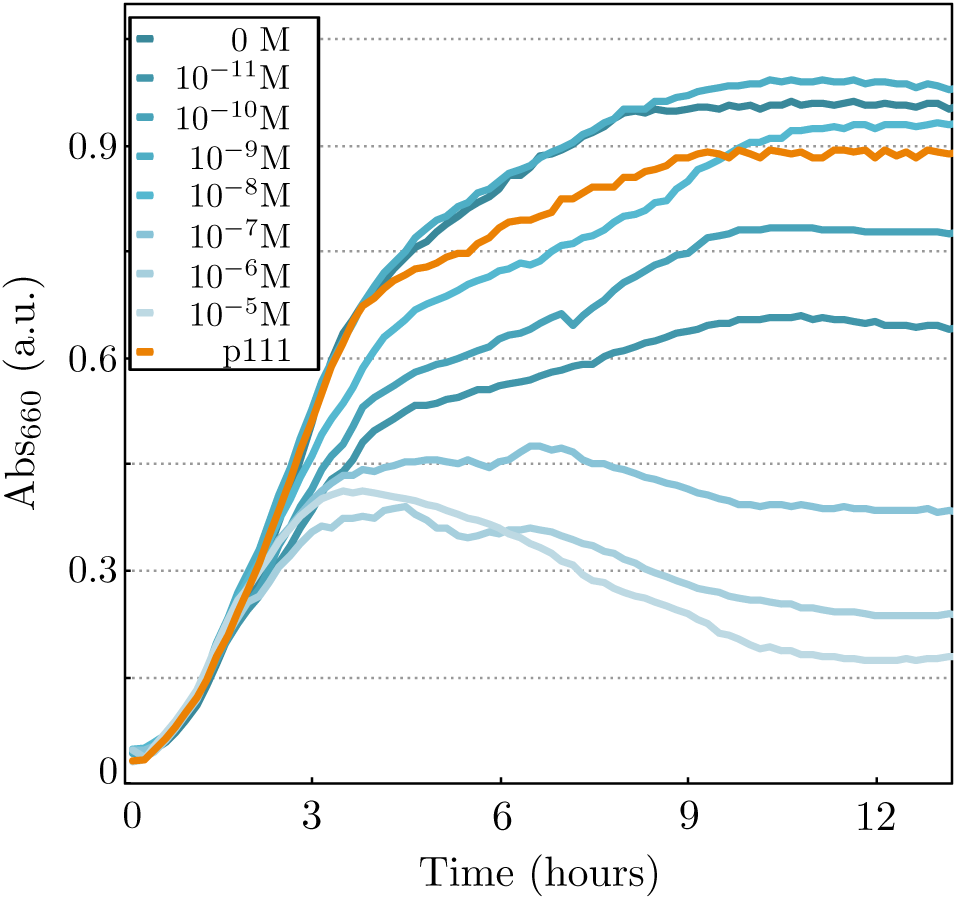
Bacterial growth is influenced by lactone concentration. Characterization of the effect of *MinC* and *JunA* expression on the growth of synthetic cells. For different concentrations of the quorum sensing molecule, the construct p011, har-boring constituive *LuxR* and conditional expresion of *MinC* and *JunA*, shows different growth rates and carrying capacity (shades of blue, average of two independent replicates). As a control p111, capable of endogenously synthesizing lactone, displays a growth rate similar to 10^−9^M of the previous case (orange).

This necessarily implies that both a decrease in growth speed and a cell death response are mediated by *MinC* and *JunA*, suggesting that this mechanism might be responsible for an active inhibition between branches and local activation associated to cellular division.

## III. DISCUSSION

Turing’s initial proposal of a diffusion-driven pattern formation mechanisms was nothing short of revolutionary for the area of developmental biology. It kickstarted a decades long search for the interaction motifs and the molecular entities that might operate this mechanism. This strategy found wide success first in chemical systems [15–18], where chemical Turing patterns were reported and studied long before the biological domain. The molecular evidence for Turing-type mechanism in biological systems has been sparse up until very recently [29–31], casting doubts on the universality of diffusion-driven instabilities. Here we have taken a different approach to analyzing naturally occurring periodic patterns, constructing *de novo* using synthetic biology components a system capable of producing periodic structures. This perspective has the added benefits of testing preconceived knowledge on how a system must operate. In this case we challenged the idea of the activator (bacteria) being a immutable element in terms of diffusible properties as the process unfolded. With the expression of an elongating factor and an adhesion protein, the activator compartment actually changes its properties depending on the inhibitor density (HSL). This relaxation of the operating conditions might help expand research programs on Turing-type pattern formation, especially now that diffusion-driven mechanisms have been shown to be common but not robust in simulated transcription factor networks [67, 68], the parameter space supporting them much smaller than previously thought.

The presence of a cell consortium (resulting from HSL-induced differentiation), the identification of a cell population (not a molecular signal) as an activator, the facilitative differentiation effect played by the inhibitor and the existence of physical effects derived from the elongated nature of *y* cells are specific components of our synthetic design that depart from the simple AIM design depicted in figure 1a. However, the basic logic remains the same, since the facilitation of differentiation *x → y* by the HSL, effectively inhibits the growth rate of the HSL-producing *x* component of the consortium. However, given the special cell morphology of the differentiated strain, further work should help understand how these phenotypic traits modulate the spatial patterning [69].

A separate aspect tackled in this study is how embodiment affects the interactions between the agents in the system. With the expression of MinC through the presence of the signalling molecule HSL, bacteria become elongated and do not divide as often. This has a synergistic effect in the surface presentation of the adhesion protein JunA, enhancing the capacity of cells to attach with one another as well as create cohesive bundles (as shown in figure 4). These appear very soon during colony development, and their orientation seems to be the cause of directed flow/growth at the edge of the colony and the branching process. This level of detail in synthetic biology applications is often overlooked for the more simplistic and tractable computational perspective [48]. However, an embodied perspective on synthetic pattern formation has plenty of theory to draw upon [1] and can try to achieve symmetry breaking with the help of physical processes, not despite their existence. This in turn, might drive the focus of synthetic biology from a computational perspective to a functional one, with tool-kits developed to interface with known physical processes through the embodiment of the agents.

## IV. MATERIALS AND METHODS

### 1. DNA constructs and plasmids

Final genetic constructs used in this work were generated using the standard biobrick clonning techniques and enzymes: EcoRI/XbaI/SpeI/PstI restrictases and T4 DNA ligase (New England Biolabs, USA). Some DNA sequences were provided by the iGEM 2010 spring collection, including *LuxR* (C0062), *LuxI* (C0161), *pLux* (R0062), *MinC* (K299806), *GFP* (E0040), constitutive promoter (J23100), bidirectional terminator (B0014) and RBSs (B0033, B0034). *JunA* was formatted to biobrick standard 10 from a coding sequence kindly provided by L.A. Fernández. The inefficient Lux promoter *pL40* was created *de novo* by primer hybridization (Sigma Aldrich, USA). See supplementary materials for sequences of all used DNA pieces.

Genetic devices were split between two plasmids pSB1AC3 and pSB3K5, also obtained from the iGEM 2010 distribution, with high and high-intermediate copy numbers respectively. The two plasmids harbor different origins of replication, and can coexist inside a single cell. All final constructs were sequenced by the PRBB core facilities.

### 2. Bacterial strains and growth conditions

Cloning procedures were carried out in *E. coli* Top10 strain (Invitrogen, USA). Final essays were performed in *E. coli* UT5600 kindly provided by L.A. Fernández.

Colony essays were performed as follows: UT5600 cells harboring each device were fresh plated overnight from a glycerinate stored at −80C, a single colony was then grown in Lisogenic Broth (Sigma Aldrich, USA) supplemented with Chloramphenicol and Kanamycin (Sigma Aldrich, USA) for 5 hours and diluted to Abs_660_ = 0.2. A small volume (2 *µ*L) of the density adjusted cultures was dropped in the center of 5.5 cm petri dishes, filled with 5.5 mL of LB Eiken agar (Eiken Chemical, Japan) at 0.4% w/v again supplemented with Chloramphenicol and Kanamycin and, when necessary, 10^−8^ M N-[*β*-ketocaproyl]-L-homoserine lactone (Cayman Chemical Company, USA). Inoculated plates were dried for 5 minutes and grown 14h at 37C, then stored at 22C for 7 days, were data capture took place.

### 3. Data capture and processing

Assessment of lactone concentration impact on strain growth was carried out in Synergy MX microplate reader (BioTek Instruments, USA), similarly to our previously described protocol [70]. Photographs of colony pattern were taken daily with a Canon EOS with diffuse illumination. Initial and final state of the pattern formation process were captured by bright field and fluorescence microscopy with a Leica DMI6000B (Leica Mycrosystems, USA). Regularities in colony boundaries were characterized with Matlab 2013b polar transformation and FFT algorithms (MathWorks, USA). All images were processed with a background subtraction and brightness adjustment.

## Acknowledgments

The authors thank the members of the Complex Systems Lab for useful discussions, specially to Arianna Bruguera for his work as laboratory technician. This study was supported by an European Research Council Advanced Grant (SYNCOM), the Botin Foundation, by Banco Santander through its Santander Universities Global Division and by a MINECO Grant FIS2015-67616-P. This work has also counted with the support of Secretaria d’Universitats i Recerca del Departament d’Economia i Coneixement de la Generalitat de Catalunya and by the Santa Fe Institute.

